# Hierarchical Machine Learning Uncovers Topological Signatures of Autophagy Regulation by Oral Bacteria in Oral Squamous Cell Carcinoma

**DOI:** 10.64898/2025.12.29.696881

**Authors:** Hamid Laitifi Navid, Mahdi Akhavan, Pooya Jalali, Amir Barzegar Behrooz, Sanaz Vakili, Rui Vitorino, Iman Beheshti, Anil Menon, Vimi S. Mutalik, Robert J Schroth, Prashen Chelikani, Saeid Ghavami

## Abstract

Oral squamous cell carcinoma (OSCC) progression has been increasingly linked to dysbiosis of the oral microbiome. We hypothesized that pathogenic versus commensal bacteria differentially rewire host autophagy networks to either promote or inhibit OSCC progression. To test this, we constructed host–bacterium autophagy interactomes from KEGG, STRING, and curated databases, identifying key network hubs (e.g., MAPK1, STAT3) via graph-theoretic metrics. We then applied a hierarchical unsupervised machine learning pipeline, combining two-stage principal component analysis with permutation testing and linear discriminant analysis (LDA), to interrogate differences in network topology. This multi-layer approach revealed a clear separation between pro-cancer (pathogenic) and anti-cancer (commensal) bacterial network signatures, with *Fusobacterium nucleatum* and *Streptococcus mitis* emerging as dominant global outliers. Pathogenic taxa activated inflammatory–metabolic autophagy signatures (e.g., NFKB1, MYC, ACACA), whereas commensals stabilized kinase–homeostasis signaling (EGFR, PTEN, HSP90AA1). Permutation testing confirmed that these network differences were highly significant and non-random (p < 0.001). We also derived a Dysbiosis Index that robustly distinguished the pro- versus anti-cancer bacterial cohorts with high predictive power. Collectively, our findings highlight oral microbiota–autophagy network topologies as potential biomarkers of OSCC dysbiosis and as novel therapeutic targets.

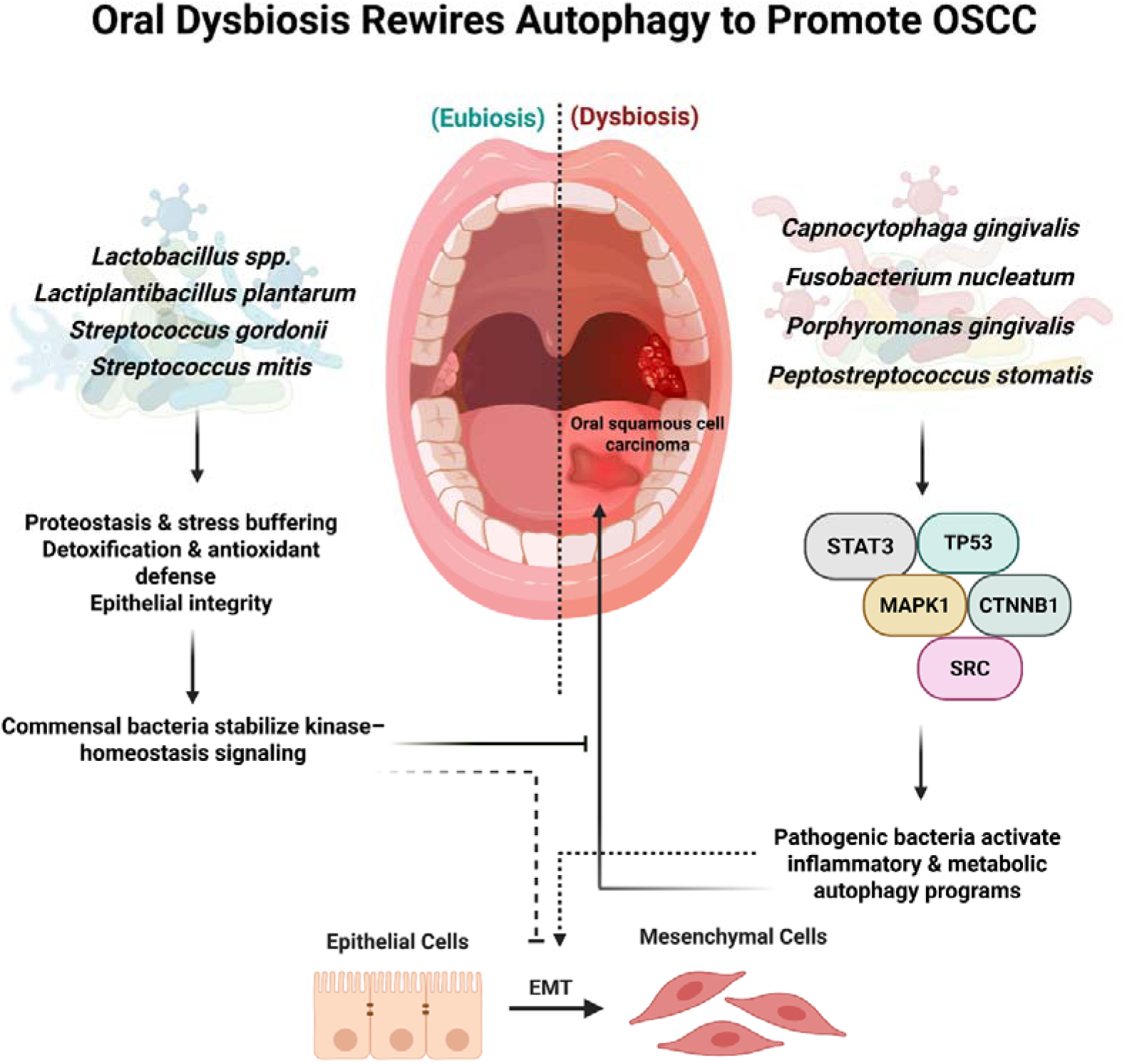

**Lay summary:** Healthy mouth bacteria help cells stay balanced and protected. When harmful bacteria take over, they disrupt cell recycling (autophagy), increase inflammation, and causing cells to become more aggressive, which can promote oral cancer development.

## Introduction

Oral squamous cell carcinoma (OSCC) is the most common malignancy of the oral cavity and remains a major clinical challenge due to poor survival rates, particularly in advanced and metastatic stages [1, 2]. The pathogenesis of OSCC extends beyond intrinsic genetic mutations to include extrinsic modulators such as chronic inflammation, bacterial dysbiosis, immune evasion, and metabolic reprogramming. These collectively promote epithelial–mesenchymal transition (EMT), autophagy deregulation, and tumor progression [3].

Evidence is emerging that the oral bacterium is not a passive bystander but as an active contributor to OSCC development [3]. Pathogenic species such as *Capnocytophaga gingivalis, Fusobacterium nucleatum*, and *Porphyromonas gingivalis* have been linked to higher tumor burden, increased invasiveness, and worse clinical prognosis. These bacteria modulate oncogenic signaling cascades including Extracellular signal-regulated kinase 1/2 (ERK1/2), signal transducer and activator of transcription 3 (STAT3), nuclear factor kappa-light-chain-enhancer of activated B cells (NF-κB), and p38 mitogen-activated protein kinase (p38 MAPK), induce matrix metalloproteinases, and hijack autophagy to facilitate tumor adaptation and survival under stress [4, 5]. Systems-level interactome analyses further suggest these species drive immune escape and therapy resistance by reprogramming host regulatory circuits [6].

In contrast, the healthy oral bacterium—comprising species such as *Lactobacillus plantarum*, *Streptococcus mitis*, and *Streptococcus gordonii*—play a cytoprotective role in epithelial homeostasis. These commensals reinforce the mucosal barrier, detoxify carcinogens, mitigate inflammation, and induce regulated autophagy, thereby reducing OSCC risk [7, 8]. Disruption of this microbial balance, however, contributes to disease onset and progression [9].

Autophagy itself is a highly conserved catabolic process that maintains proteostasis, regulates immune signaling, and governs the epithelial response to stress [10]. In the context of cancer, autophagy displays a dual role: while cytoprotective in early disease, it may switch to a tumor-promoting phenotype during later stages or in response to therapy-induced stress [11]. Oral bacterial products such as Pathogen-associated molecular patterns (PAMPs) and short-chain fatty acids (SCFAs)., and cytokines modulate this pathway by engaging mechanistic target of rapamycin (mTOR), STAT3, and MAPK signaling cascades, resulting in context-dependent autophagy regulation [12, 13].

Despite growing recognition of this microbe-autophagy-tumor axis, few studies have systematically compared the impact of multiple bacterial species on host autophagy and network reprogramming in OSCC [14]. Existing models largely focus on individual pathogens and lack integrative mapping of divergent bacterial strategies across a shared host network landscape [15].

To address this gap, we developed an integrative systems-level framework to interrogate how oral bacteria reprogram autophagy-centered host signaling in OSCC. We specifically tested the hypothesis that pathogenic and commensal oral bacteria employ fundamentally distinct network-level strategies—either reinforcing inflammatory and pro-survival signaling or preserving epithelial homeostasis—through differential rewiring of autophagy-associated regulatory hubs.

## 2. Materials and Methods

To test this hypothesis, we employed a multi-layered systems biology and unsupervised machine learning approach. Using the Kyoto Encyclopedia of Genes and Genomes (KEGG) and curated host–microbe interactome databases, we constructed an integrated map of autophagy-related signaling hubs influenced by OSCC-associated oral bacteria. We then applied a hierarchical unsupervised analytical pipeline to characterize network topological perturbations induced by ten representative oral bacterial species. Principal Component Analysis (PCA) was used to identify dominant and latent patterns across bacterial cohorts, followed by permutation testing, discriminant analysis, and pathway-level scoring to resolve functionally distinct bacterial signatures.

### 2.1 Identification of OSCC-Associated Host Proteins

To identify key proteins associated with oral squamous cell carcinoma (OSCC), the STRING database (Version 12.0; https://string-db.org/) was queried. STRING is a comprehensive resource that integrates known and predicted protein–protein interactions (PPIs) derived from diverse evidence streams, including high-throughput experimental datasets, curated pathway knowledgebases, gene co-expression patterns, computational predictions (e.g., genomic context and text mining), and orthology-based information transfer. Using OSCC-related disease terms and pathway descriptors, **125 proteins** implicated in OSCC biology or associated signalling networks were retrieved (Supplementary Table 1). These proteins served as the foundational OSCC-related host gene set for downstream comparative and integrative analyses.

### 2.2 Compilation of Autophagy Signaling Genes

To establish a comprehensive reference framework for autophagy-associated signalling, genes involved in autophagy were compiled by interrogating multiple curated databases, including STRING (https://string-db.org/), Reactome (https://reactome.org/), UniProt (https://www.uniprot.org/), KEGG (https://www.genome.jp/kegg/), and HMDB (https://www.hmdb.ca/). Pathway identifiers, GO terms, and accession numbers covering macroautophagy, selective autophagy, autophagosome assembly, lysosomal pathways, and phagophore formation were collected as shown in Table 1. All gene lists were merged and deduplicated using official HGNC symbols as unique identifiers. This process yielded a consolidated set of **1,018 unique autophagy-related genes** (Supplementary Table 2), which formed the canonical autophagy network for subsequent integration with OSCC and bacteria-derived host interactomes.

**Table 1.** Pathway identifiers and database accession numbers used for systematic retrieval of autophagy-related genes.

| STRING |  |  |  |
| --- | --- | --- | --- |
| GO:0006914 | Autophagy | GO:0044754 | Autolysosomes |
| GO:0016236 | Macroautophagy | GOCC:0061908 | Phagophore |
| GO:0044804 | Autophagy of the Nucleus | GO:0000407 | Phagophore Assembly Site |
| GO:0005776 | Autophagosome | GO:0034497 | Protein Localization to Phagophore Assembly Site |
| GO:0061912 | Selective Autophagy | GO:0034045 | Phagophore assembly site membran |
| GO:0000045 | Autophagosome Assembly | GO:0097632 | Extrinsic component of phagophore |
| GOCC:0005776 | Autophagosome | GOCC:0097632 | Phagophore |
| **Reactome** |  | **KEGG** |  |
| R-HSA-9612973 | Autophagy | hsa04140 | Autophagy (or macroautophagy) |
| R-HSA-1632852 | Macroautophagy | hsa04136 | Autophagy - other - Homo sapiens (human) |
| **UNIPROT** |  |  |  |
| KW-0072 | Autophagy |  |  |

### 2.3 Defining Host Targets of OSCC-Relevant Oral Bacteria

To characterize the host proteins targeted by oral bacterial taxa implicated in OSCC, we compiled experimentally supported host–bacterium interaction data from peer-reviewed literature. Ten oral bacterial species were selected based on clinical relevance and experimental evidence linking them either to OSCC progression (e.g., *Fusobacterium nucleatum*, *Porphyromonas gingivalis*, *Capnocytophaga gingivalis*, *Prevotella stomatis*, *Prevotella intermedia*) or to maintenance of oral epithelial homeostasis (e.g., *Streptococcus mitis*, *Lactobacillus plantarum*, *Lactobacillus sp.*, *Streptococcus gordonii*, *Streptococcus anginosus*) (Table 2).

**Table 2:** Host-Associated Oral Bacteria in OSCC: Cancer Context and Sequencing Approaches.

| Bacteria | Cancer type (OSCC, HNSCC etc) | Sequencing methods (16S rRNA, Shotgun, RNA seq) | Reference |
| --- | --- | --- | --- |
| *Fusobacterium nucleatum* | OSCC (oral cavity OSCC tissue) | 16S rRNA gene sequencing of tumor vs control oral tissues | [16, 17] |
| *Porphyromonas gingivalis* | OSCC | RNA-seq (SCC-25 oral squamous carcinoma cells stimulated with *P. gingivalis* W83 membrane components) | [18] |
| *Capnocytophaga gingivalis* | OSCC (saliva and tissue) | 16S rRNA gene sequencing of saliva microbiome in OSCC vs controls; FISH in OSCC tissues | [19] |
| *Prevotella intermedia* | OSCC (saliva OSCC cohort including periodontal pathogens) | 16S rRNA gene sequencing of saliva microbiome in OSCC vs controls (Reporting *Prevotella* spp., including *P. intermedia*) | [19] |
| *Prevotella stomatis* | OSCC (tumor and saliva microbiome) | 16S rRNA gene sequencing across OSCC clinical samples (Tumor, Saliva, Mucosal swabs) | [20] |
| *Streptococcus mitis* | HNSCC including oral cavity SCC | 16S rRNA gene sequencing of OSCC and control tissues, reporting *Streptococcus* spp. composition | [16, 21] |
| *Streptococcus gordonii* | OSCC (tumor microbiome) | 16S rRNA sequencing of OSCC tumor vs non-tumor tissue | [16, 22] |
| *Streptococcus anginosus* | OSCC / HNSCC (tissue-based microbiome) | 16S rRNA gene sequencing and targeted detection in OSCC tissue | [16, 23] |
| *Lactobacillus plantarum* | OSCC (saliva) | 16S rRNA gene sequencing of saliva (Reporting increased *Lactobacillus* in OSCC patients) | [19] |
| *Lactobacillus* spp. (other than *L. plantarum*) | OSCC (saliva and tissue) | 16S rRNA sequencing of saliva and OSCC tissue microbiome | [19] |

**Table 3.**
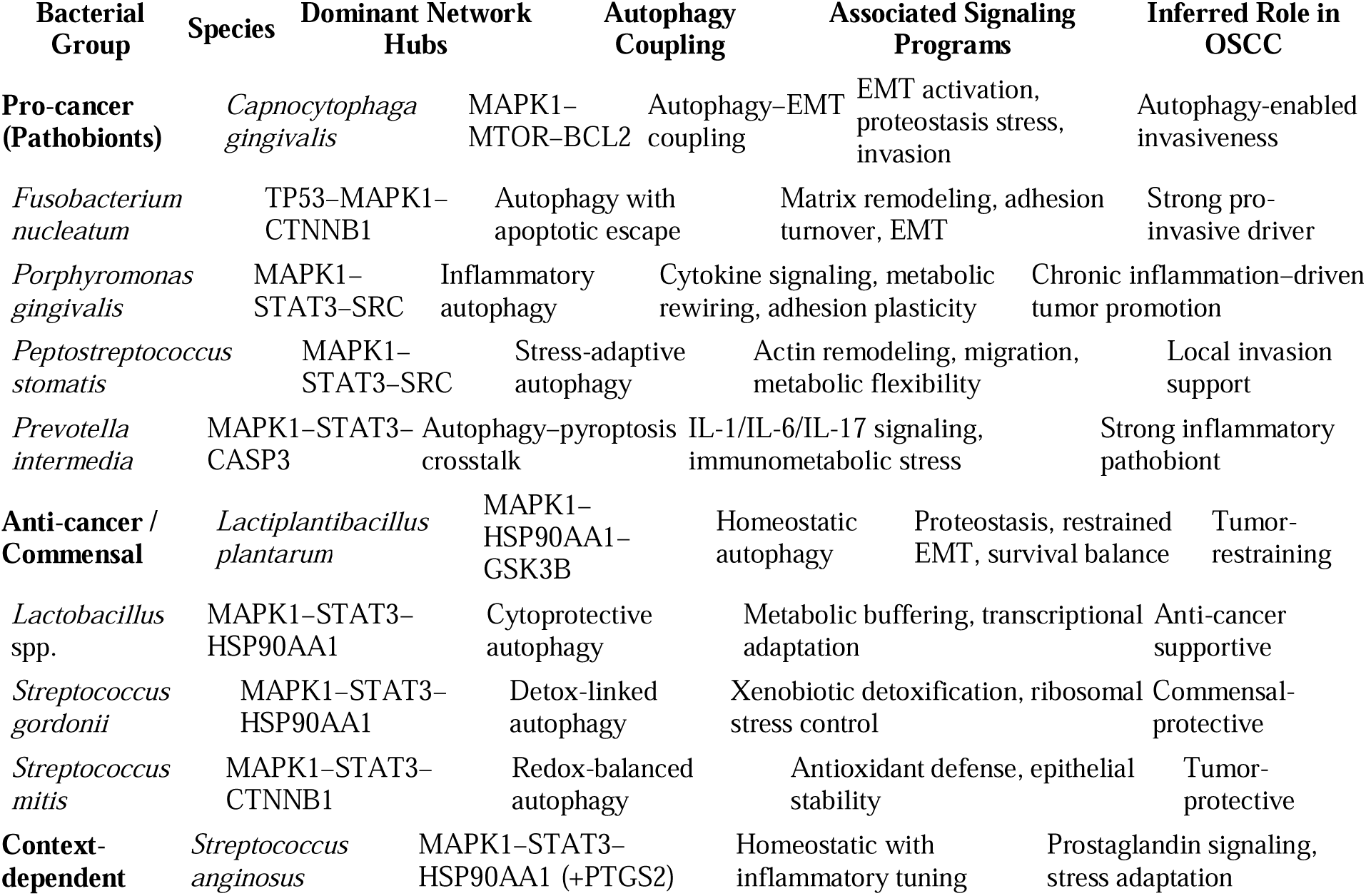
Functional classification of oral bacteria based on autophagy-centered signalling topology and inferred relevance to OSCC biology.

In addition, the clinical context, sequencing methodologies based on the indexed studies supporting the inclusion of each of these ten taxa are summarized in Table 2, providing an explicit link between each bacterium and OSCC or OSCC-relevant host responses.

For each bacterium, the curated host target list was integrated with both the OSCC gene set (Section 2.1) and the autophagy gene set (Section 2.2) to generate bacterium-specific candidate host signalling modules. These combined gene lists were then used to construct bacterium-specific first-order human PPI networks through NetworkAnalyst, using STRING v11.0 with a high-confidence cut-off (>0.7). Within each interactome, Degree Centrality and Betweenness Centrality metrics were applied to highlight influential hub nodes.

By jointly analyzing (i) OSCC-derived hub proteins, (ii) autophagy-related hub proteins, and (iii) literature-derived bacterial host targets, a set of ten bacteria-centric host protein modules was obtained. Taxon-specific interactomes were then generated for each bacterial species and visualized using the Steiner Forest algorithm, which enabled extraction of the primary structural backbone and key regulatory axes for each network (Supplementary Tables 3–22).

### 2.4 Construction of Bacterium-Specific Host PPI Networks

Each bacterial species yielded a distinct high-confidence human PPI subnetwork representing its predicted influence on host signalling. Degree centrality was calculated for each node as a measure of local regulatory influence:

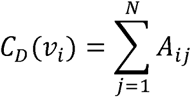

*A_ij_* represents the element of the adjacency matrix indicating whether a connection exists from node *i* to node *j*.. These centrality values were used to build numerical representations of each bacterial interactome.

### 2.5 Feature Matrix Generation and Normalization

All ten interactomes were integrated into a unified **10 × N** feature matrix, where each row corresponded to a bacterial species and each column to a unique host gene identified across networks. Missing genes (i.e., genes absent from a given bacterial network) were imputed as zero, representing true topological absence rather than missing data.

Z-score normalization was employed to standardize each feature prior to analysis. For a given feature *j*, the normalized value for observation *i* was computed as: 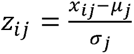 where *x_ij_* denotes the original measurement of feature *j* for observation *i, µ_j_* is the mean of feature *j* across all observations, and *σ_j_* is the corresponding standard deviation. This strategy ensured that downstream machine learning analyses captured biological structure rather than differences in absolute scale.

### 2.6 Hierarchical Unsupervised Machine Learning

To analyze the high-dimensional topological landscape of bacterial host perturbation, we employed a two-stage PCA technique.

1. **Global PCA** was first applied to all ten bacterial profiles to identify the dominant axes of variance. This analysis consistently highlighted *F. nucleatum* (pathogenic) and *S. mitis* (commensal) as global outliers due to their unusually broad or unusually focused signaling footprints.
2. **Zoomed-In PCA** was then performed after temporarily removing these two species to resolve subtler patterns within the remaining core bacterium (N = 8). This hierarchical design allowed us to uncover both macro-level (global) and micro-level (cohort-specific) topological signatures.

### 2.7 Statistical Validation

To determine whether the observed PCA structure reflected non-random biological organization, we performed a permutation-based Monte Carlo test. All features were standardized (z-score normalization). For each of *B* = 1,000 replicates, entries within each feature column were permuted independently to disrupt inter-feature correlations while preserving marginal distributions. PCA was recomputed for each permuted matrix, and the test statistic *T*(leading eigenvalue, *λ*_1_; cumulative variance of the first *k* components was also examined) was recorded. The empirical p-value was calculated as:

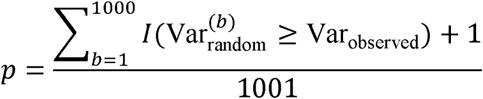

A p-value < 0.05 supported the presence of non-random, biologically meaningful structure.

To evaluate pathway-level specificity, Fisher’s Exact Test was applied to assess enrichment of high-centrality genes (e.g., inflammatory, metabolic, homeostasis-kinase signatures) in pathogenic versus commensal cohorts.

To test cohort separability, Linear Discriminant Analysis (LDA) was performed on PCA-transformed data. A composite Dysbiosis Index, defined as the difference between normalized Inflammation and Homeostasis Scores, provided a quantitative metric summarizing bacterial pathogenic potential.

### 2.8 Overall Analytical Framework

By integrating OSCC signalling, autophagy pathway biology, literature-derived bacterial targets, human interactomes, graph-theoretic structure, hierarchical PCA, and non-parametric statistical validation, this framework enabled a systems-level reconstruction of how pathogenic and commensal oral bacteria differentially reprogram host signalling in OSCC.

### 2.9 Independent Clinical and Biological Validation

To assess the translational potential of computationally predicted autophagy-related hubs, a dual-layered validation was done using external clinical data sets. Initially, a transcriptome profiling was performed using GEO data set GSE37991, which included tumor and non-cancer biopsies for 40 OSCC patients was selected. Differential expression was done using the Wilcoxon signed-rank test. Genes were assessed for their topological significance in our host-microbe interactome; a list was considered significant for differential expression (DEGs) if satisfying p-values < 0.05 and |log2 fold change| > 1. Secondly, to validate these results at a translational level, expression data for proteins under investigation was queried using Human Protein Atlas (HPA) database [(https://www.proteinatlas.org)] for IHC data which was used to determine intensity (High, Medium, Low, or Not Detected) differences in tumor biopsies and normal oral mucosa, while accounting for the appearance of clinical manifestations of predicted pathogenic and homeostatic expression signatures.

The workflow has summarized in Scheme 1.

**Scheme 1.**
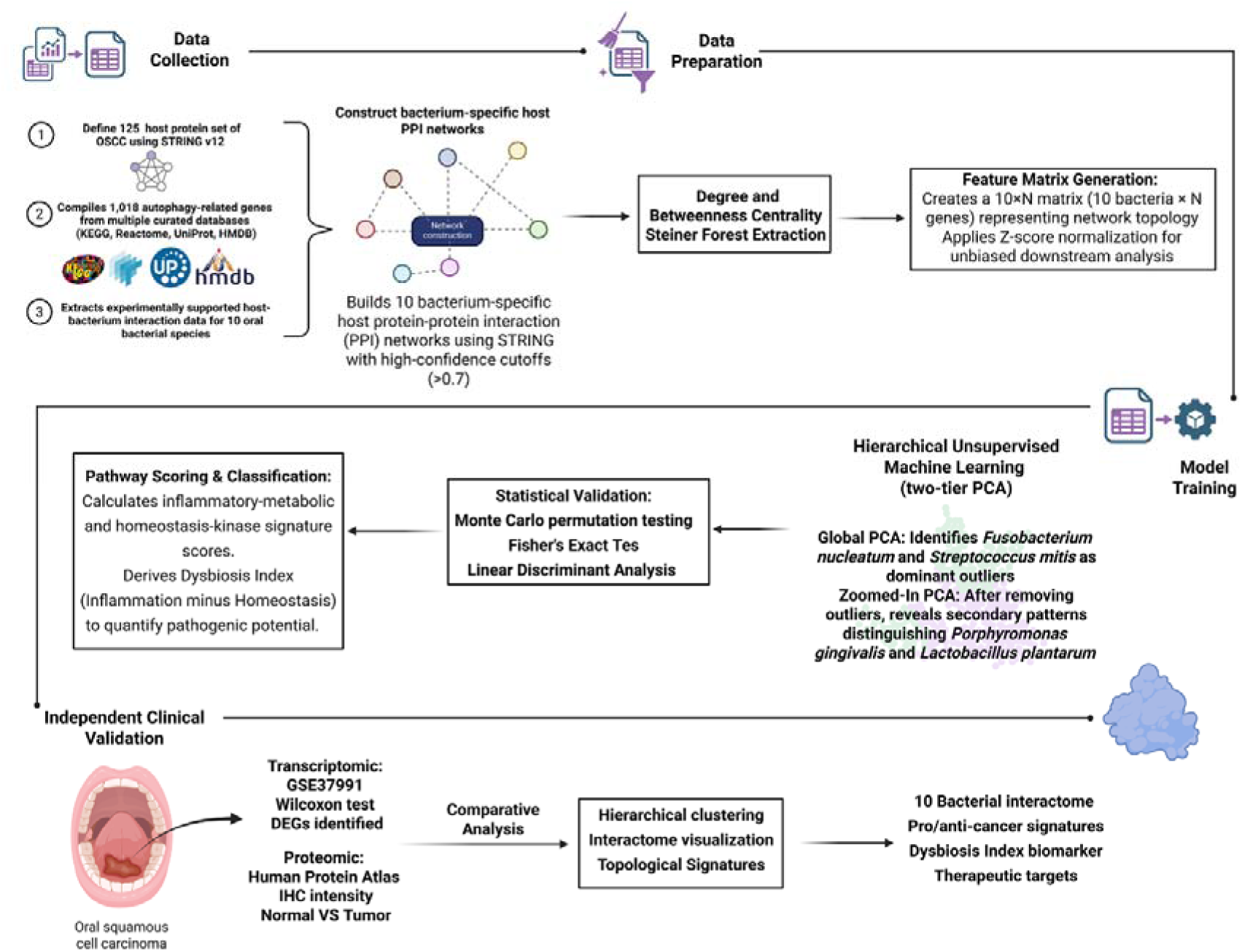
Integrated analytical workflow of the study. Curated host–bacteria interaction data and autophagy-related gene sets were assembled to construct bacterium-specific host protein–protein interaction networks. Network topology features were extracted and organized into a unified feature matrix. Hierarchical unsupervised machine-learning analyses identified dominant and secondary signalling signatures, followed by statistical validation. Independent transcriptomic and proteomic datasets were used for clinical validation, enabling classification of oral bacteria into pro-cancer or homeostatic groups and identification of dysbiosis-associated therapeutic targets.

## Result

### 1. Pro-cancer oral bacteria converge on inflammatory and EMT-permissive autophagy networks in OSCC

To clarify shared oncogenic signaling principles, we grouped oral bacteria associated with tumor promotion—*Capnocytophaga gingivalis* (Figure 1A), *Fusobacterium nucleatum* (Figure 1B), *Porphyromonas gingivalis* (Figure 1C), *Peptostreptococcus stomatis* (Figure 1D), and *Prevotella intermedia* (Figure 1F)—into a unified interactome analysis. Despite species-specific features, all pathobionts converged on MAPK1- and STAT3-centered signaling hubs that coupled autophagy regulation with inflammatory signaling, epithelial–mesenchymal transition (EMT), cytoskeletal remodeling, and metabolic rewiring.

**Figure 1.**
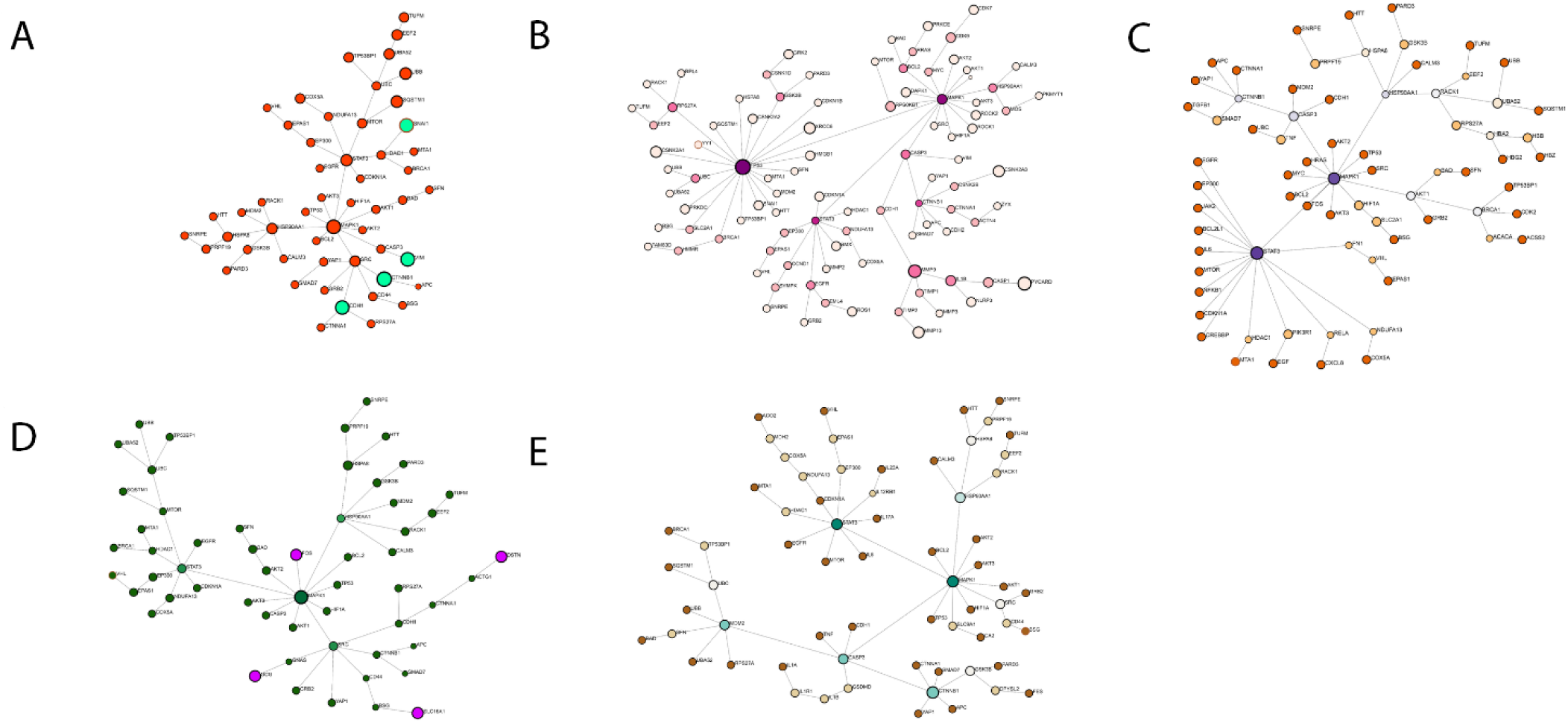
Pro-cancer oral bacteria converge on inflammatory autophagy–EMT signaling networks in OSCC. Integrated protein–protein interaction network summarizing signaling modules induced by pro-cancer oral bacteria (*Capnocytophaga gingivalis* (A)*, Fusobacterium nucleatum* (B)*, Porphyromonas gingivalis* (C)*, Peptostreptococcus stomatis* (D), and *Prevotella intermedia* (E)). The network highlights dominant hubs centered on MAPK1, STAT3, TP53, SRC, and CASP3, linking autophagy regulators (mTOR, BCL2, SQSTM1) with EMT drivers (CTNNB1, CDH1, VIM), inflammatory cytokines (IL1, IL6, IL17, IL23), matrix remodeling enzymes (MMPs), and cytoskeletal effectors. Node size reflects degree centrality and edge thickness denotes interaction confidence. Color gradients distinguish autophagy, inflammatory, EMT, apoptotic, and metabolic modules. The composite topology illustrates how pathobionts promote inflammatory autophagy, apoptotic tolerance, and invasive epithelial plasticity in OSCC.

*C. gingivalis* established a MAPK1–mTOR–B-cell lymphoma 2 (BCL2) axis linking autophagy flux to EMT drivers (Snail family transcriptional repressor 1 (SNAI1), catenin beta-1 (CTNNB1), and vimentin (VIM)), suggesting autophagy-enabled invasiveness. *F. nucleatum* formed a Tumor protein p53 (TP53)–MAPK1–CTNNB1 tri-hub, synchronizing apoptotic escape, matrix remodeling (MMP2/9/13), and adhesion turnover. *P. gingivalis* and *P. stomatis* shared a MAPK1–STAT3–Proto-oncogene tyrosine-protein kinase Src (SRC) architecture, integrating cytokine-driven transcription, actin dynamics (DSTN, ACTG1), and metabolic shuttling (SLC16A1), consistent with inflammatory stress adaptation and migration. Prevotella intermedia uniquely extended this framework by coupling Caspase-3 (CASP3), gasdermin D (GSDMD), interleukin-1 (IL-1), interleukin-6 (IL-6), interleukin-17 (IL-17), and interleukin-23 (IL-23) signaling, linking autophagy to pyroptosis and immunometabolic inflammation.

Collectively, pro-cancer bacteria reshape epithelial signaling toward inflammatory autophagy, apoptotic tolerance, and EMT-driven plasticity, defining a pathobiont gradient in which autophagy supports invasion rather than homeostasis (Figure 1).

### 2. Commensal and anti-cancer oral bacteria stabilize homeostatic autophagy and epithelial integrity in OSCC

In contrast, *Lactiplantibacillus plantarum* (2A), *Lactobacillus* (2B), Streptococcus gordonii (2C), *Streptococcus mitis* (2D), and the mixed responder *Streptococcus anginosus* (2E) formed a second functional group characterized by balanced, cytoprotective autophagy networks. These taxa consistently centered on MAPK1–STAT3–proteostasis hubs, but with restrained EMT engagement and prominent antioxidant, detoxification, and metabolic buffering modules.

*L. plantarum* and Lactobacillus organized MAPK1, heat shock protein 90 alpha family class A member 1 (HSP90AA1), glycogen synthase kinase-3 beta (GSK3B), STAT3 triads, reinforcing chaperone-guided proteostasis and controlled survival signaling while dampening EMT. *S. gordonii* emphasized detoxification (Glutathione S-transferase pi 1 (GSTP1) and cytochrome P450 family 2 subfamily E member 1 (CYP2E1)) and ribosomal stress control, aligning autophagy with translational fidelity. S. mitis extended this architecture to include oxidative stress defense (Catalase (CAT) and superoxide dismutase 1 (SOD1)) and xenobiotic metabolism (Cytochrome P450 (CYP) and aldehyde dehydrogenase (ALDH) enzyme families), preserving epithelial stability via CTNNB1-mediated junctional control. *Streptococcus anginosus* retained this homeostatic backbone but added a Prostaglandin-endoperoxide synthase 2 (PTGS2)-dependent prostaglandin arm, indicating context-dependent inflammatory tuning without full oncogenic conversion.

Together, commensal bacteria bias autophagy toward stress resolution, redox balance, and epithelial maintenance, opposing the inflammatory and invasive signaling imposed by pathobionts (Figure 2).

**Figure 2.**
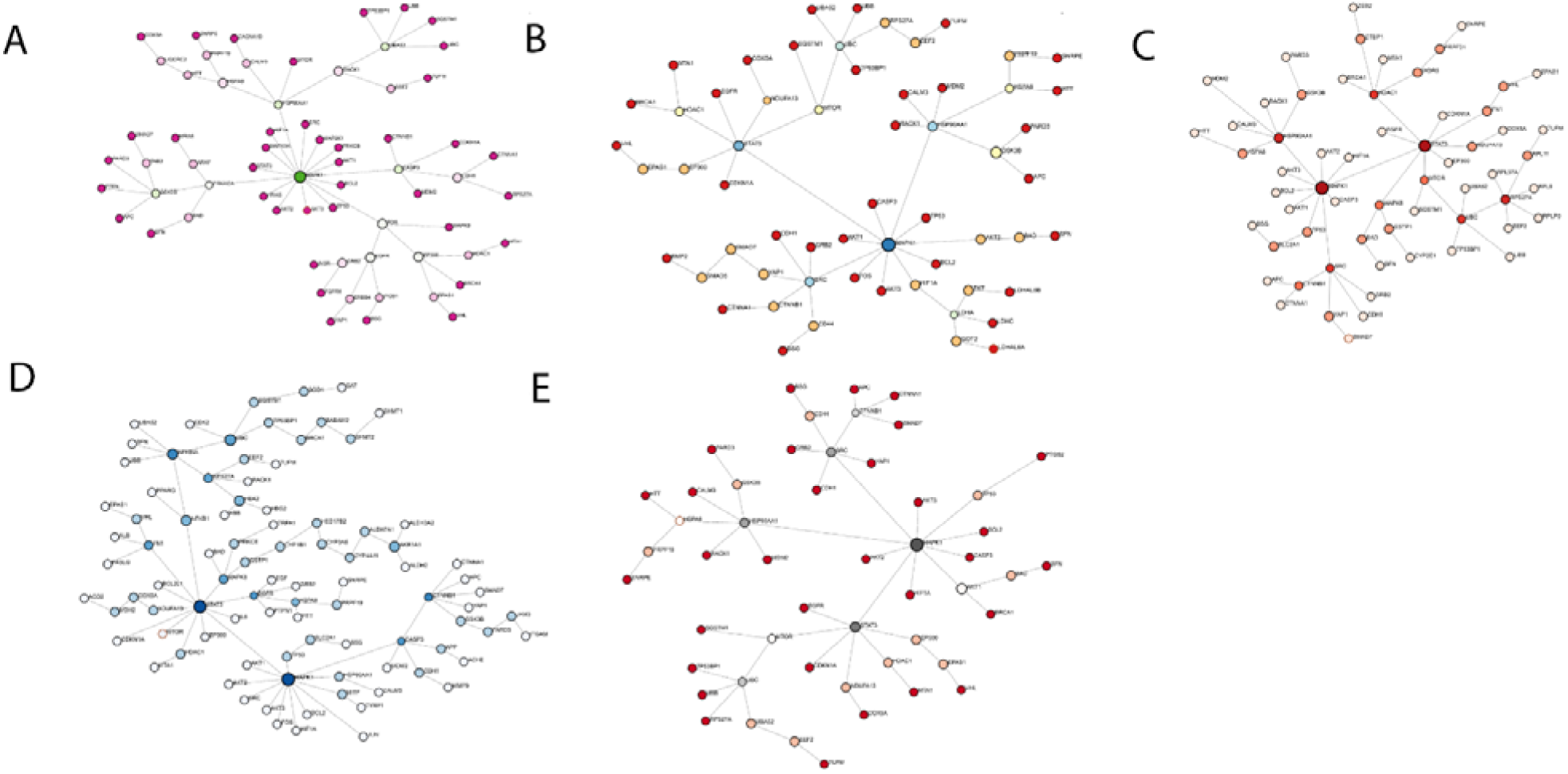
Commensal oral bacteria promote homeostatic autophagy, proteostasis, and epithelial stability in OSCC. Integrated interactome representing signaling networks induced by commensal and anti-cancer oral bacteria (*Lactiplantibacillus plantarum, Lactobacillus, Streptococcus gordonii, Streptococcus mitis,* and *Streptococcus anginosus*). The network is organized around MAPK1–STAT3–HSP90AA1/GSK3B hubs, linking autophagy regulation with proteostasis (HSPA8, CALM3), antioxidant defense (CAT, SOD1), xenobiotic detoxification (CYP and ALDH families), metabolic buffering (LDHA, TKT), and controlled epithelial adhesion (CTNNB1, CDH1). Node size and edge thickness indicate centrality and interaction confidence, respectively. The architecture reflects cytoprotective, redox-balanced autophagy that preserves epithelial integrity and limits EMT compared with pathogenic bacterial networks.

### 3. Hierarchical unsupervised learning reveals dominant and secondary topological signatures in bacterial host-interaction networks

To test whether bacterial host-interaction network topology alone can distinguish pro-cancer from anti-cancer oral bacteria, we embedded all ten bacterial gene-interaction networks into a shared high-dimensional feature space and applied unsupervised principal component analysis (PCA). This approach was deliberately label-free, allowing the model to identify intrinsic structure without prior biological assumptions.

When all ten bacterial networks were analyzed together, the PCA projection did not immediately resolve a simple binary separation. Instead, the embedding adopted an L-shaped configuration dominated by two extreme outliers (Figure 3A). *Fusobacterium nucleatum* separated strongly along the first principal component (PC1), whereas *Streptococcus mitis* was uniquely displaced along PC2. The remaining eight bacteria collapsed tightly near the origin, indicating that the strong and biologically distinct signatures of these two species overwhelmed more subtle variance contributions.

**Figure 3.**
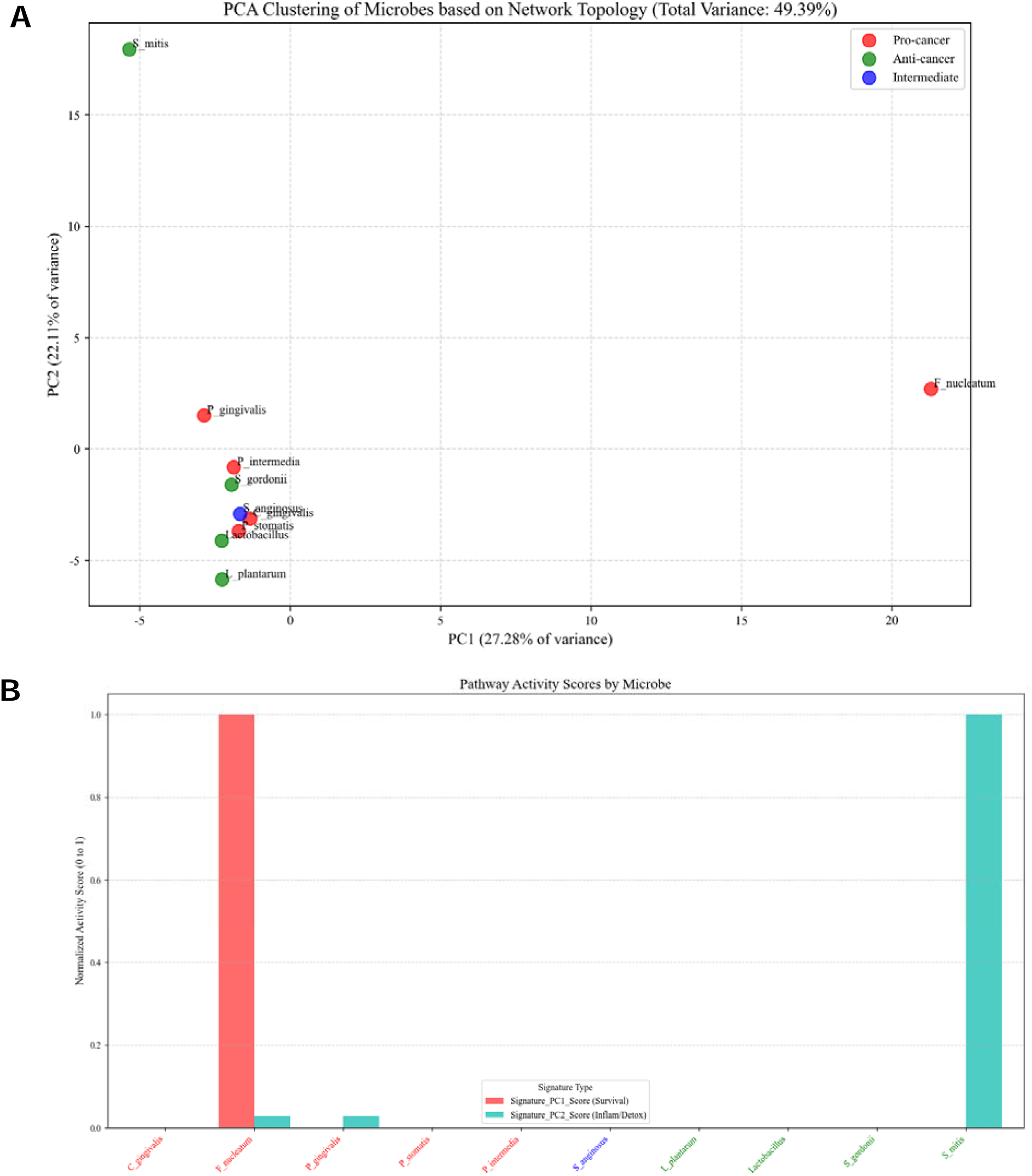
Global unsupervised PCA reveals two dominant, opposing bacterial network signatures. (A) Principal component analysis (PCA) of the full gene-network embeddings for all ten bacterial species. Each point represents one bacterial network projected onto PC1 and PC2. The projection forms an L-shaped configuration dominated by two outliers: *Fusobacterium nucleatum* (extreme along PC1) and *Streptococcus mitis* (extreme along PC2), while the remaining eight species cluster near the origin. (B) Normalized pathway-signature activity scores derived from PCA loadings. PC1 corresponds to a Survival/Cell-Cycle axis (e.g., *BCL2*, *CCND1*), maximally activated in *F. nucleatum*, whereas PC2 corresponds to a Detoxification–Autophagy axis (e.g., *ALDH* family genes, *SQSTM1*), uniquely enriched in *S. mitis*.

Pathway-level scoring confirmed this structure (Figure 3B). PC1 was enriched for survival and cell-cycle regulators (e.g., *BCL2*, *CCND1*), maximally activated in *F. nucleatum*, while PC2 mapped to a detoxification–autophagy axis involving *SQSTM1* and *ALDH* family genes, uniquely enriched in *S. mitis*. Thus, the first unsupervised decomposition revealed two dominant but opposing biological programs, insufficient to resolve the remaining species.

### 4. Removal of dominant outliers uncovers secondary mechanistic programs among remaining bacteria

To expose masked variance, we recalculated PCA after removing *F. nucleatum* and *S. mitis*. The resulting “zoomed-in” embedding again produced an L-shaped structure, now revealing *Porphyromonas gingivalis* and *Lactiplantibacillus plantarum* as secondary outliers (Figure 4A). The remaining six bacteria clustered centrally, consistent with mixed or weaker pathway engagement.

**Figure 4.**
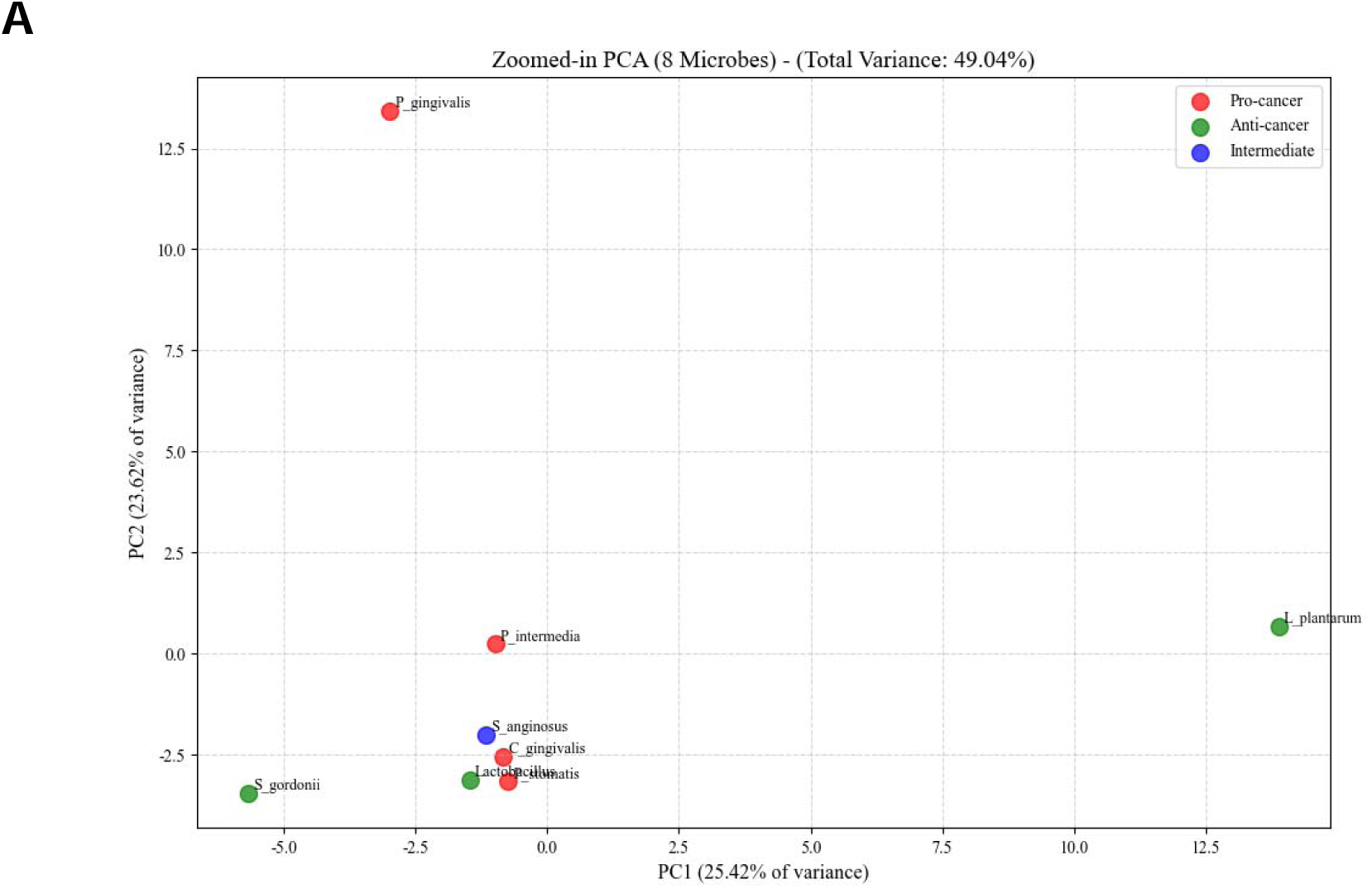
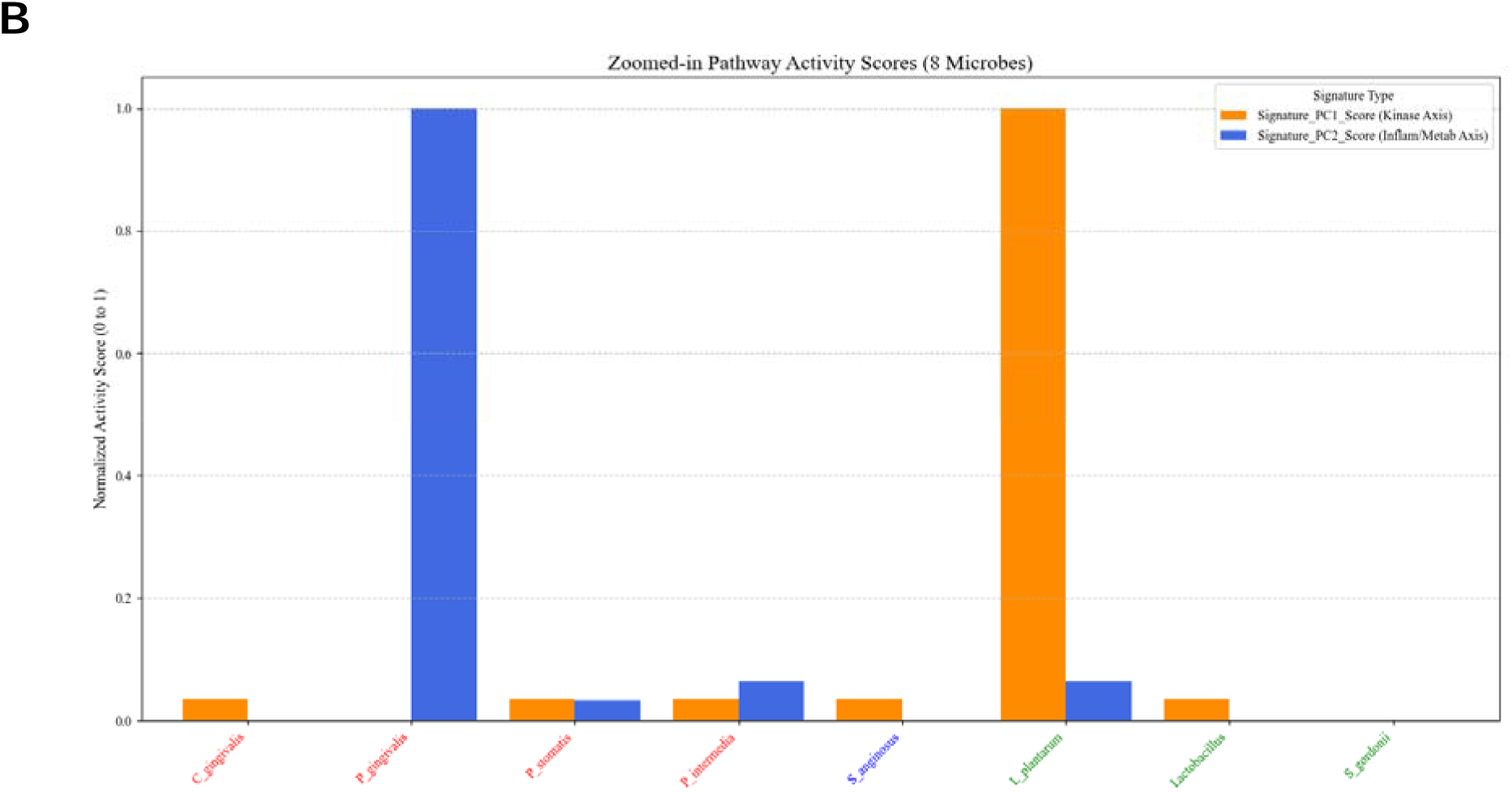
Secondary PCA resolves inflammatory versus homeostatic bacterial programs after removal of dominant outliers. (A) PCA of the eight-bacterium subset after excluding *F. nucleatum* and *S. mitis* reveals a second L-shaped configuration, with *Porphyromonas gingivalis* and *Lactiplantibacillus plantarum* emerging as orthogonal outliers and the remaining bacteria clustering centrally. (B) Corresponding pathway-activity scores defining two secondary signatures: a Regulated-Kinase/Homeostasis axis (PC1; enriched for *EGFR/PTEN*-associated modules; maximal in *L. plantarum*) and an Inflammation–Metabolic axis (PC2; enriched for *NFKB1*, *MYC*, *ACACA*; maximal in *P. gingivalis*).

Pathway scoring showed that *P. gingivalis* maximally activated an inflammatory–metabolic signature (PC2), enriched for *NFKB1*, *MYC*, and *ACACA*, consistent with its known immunometabolic pathogenicity (Figure 4B). In contrast, *L. plantarum* maximally loaded on a regulated kinase–homeostasis axis (PC1), enriched for *EGFR* and *PTEN* modulators, aligning with epithelial-

### 5. Permutation testing confirms non-random, biologically encoded network topology

To determine whether the observed variance structure reflected true biological organization rather than stochastic noise, we performed a Monte Carlo permutation test (1,000 iterations) on the full feature matrix. Randomized datasets consistently produced low total explained variance (PC1 + PC2 ≈ 28.5%), forming a tight null distribution. In contrast, the biological dataset exhibited a total explained variance of **49.4%**, falling far outside the null envelope (p < 0.001; Figure 5A).

**Figure 5.**
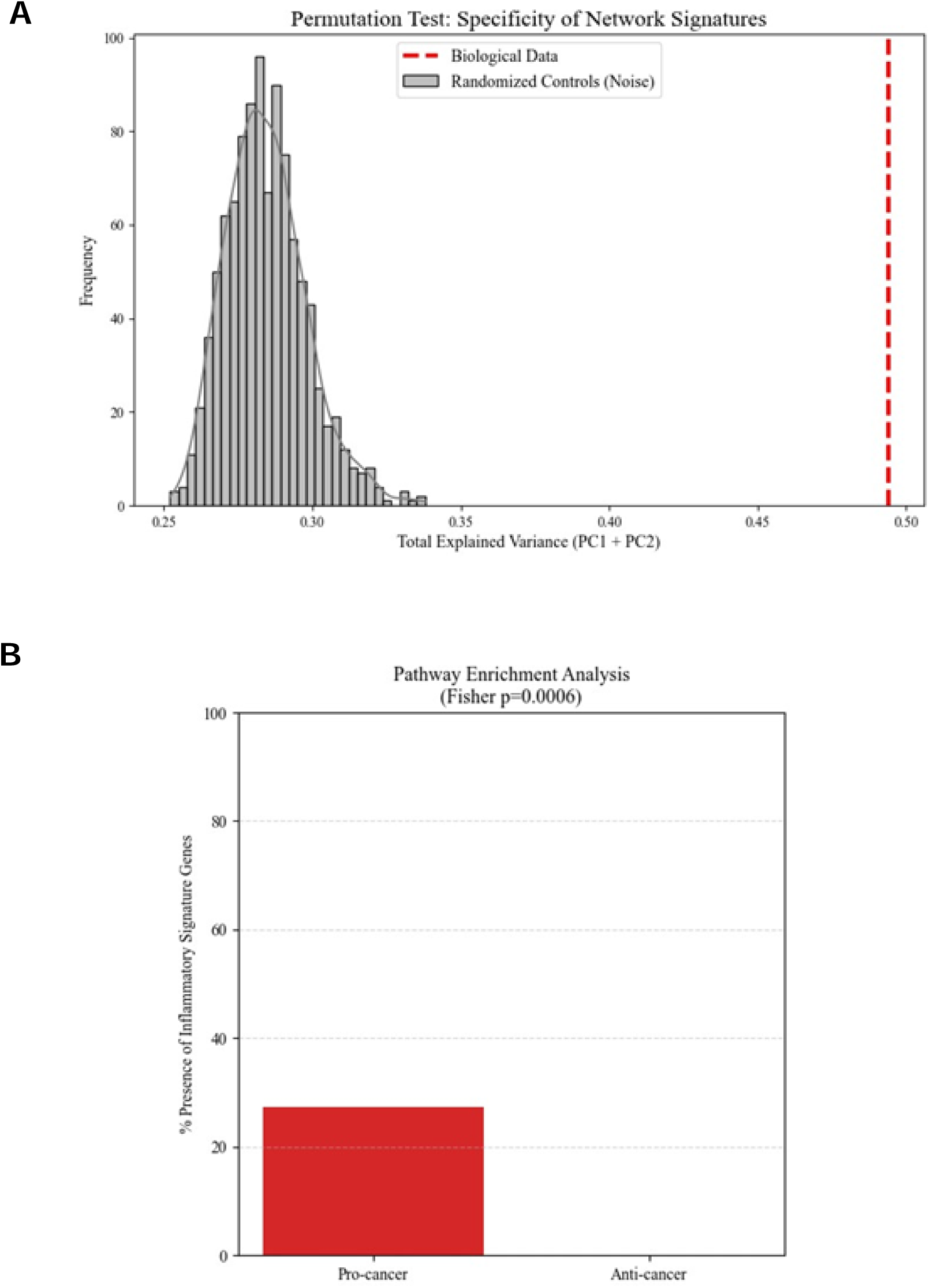
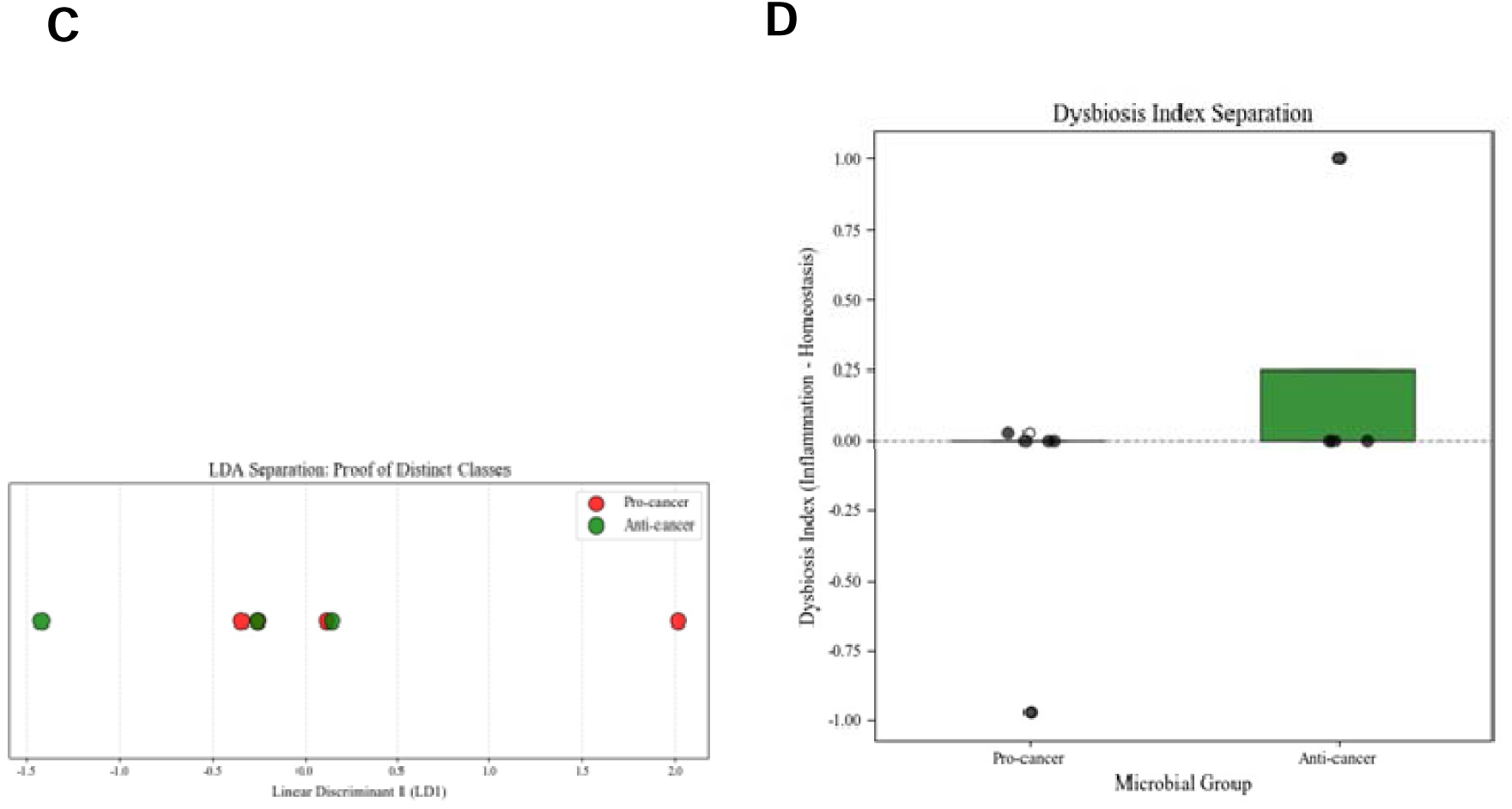
Statistical and classifier-based validation of non-random topology and cohort-level separation. (A) Monte Carlo permutation test (1,000 iterations) showing the null distribution of total explained variance (PC1 + PC2) under randomized feature matrices versus the observed biological variance (49.4%), demonstrating non-random and statistically robust topological structure (p < 0.001). (B) Fisher’s exact test demonstrating selective enrichment of a defined Inflammatory–Metabolic Signature (*NFKB1, RELA, TNF, MYC, ACACA*) in pro-cancer bacterial networks (p = 0.00057) with absence of enrichment in anti-cancer commensals. (C) Linear Discriminant Analysis (LDA) of high-dimensional network embeddings demonstrating clear linear separability between pro-cancer and anti-cancer bacterial cohorts. (D) [original Figure 17B] Composite Dysbiosis Index (Inflammation Score − Homeostasis Score) highlighting functional polarity: pro-cancer bacteria show positive dysbiosis values, whereas commensal bacteria cluster near zero or negative values.

### 6. Inflammatory–metabolic pathway enrichment and classifier-based validation define two stable bacterial classes

To directly test whether inflammatory pathway activation is selectively enriched in pro-cancer bacteria, we quantified a predefined Inflammatory–Metabolic Signature (*NFKB1, RELA, TNF, MYC, ACACA*). Fisher’s exact testing revealed significant enrichment exclusively in pro-cancer bacteria (p = 0.00057), with zero representation in anti-cancer commensals (Figure 5B).

Orthogonal validation using Linear Discriminant Analysis (LDA) demonstrated clean linear separability between pathogenic and commensal cohorts (Figure 5C). A composite Dysbiosis Index (Inflammation − Homeostasis) further amplified this polarity, with pathogenic bacteria showing strongly positive scores and commensals clustering near zero or negative values (Figure 5D. Together, these analyses provide convergent evidence that pro-cancer and anti-cancer bacteria occupy distinct and statistically stable topological regimes.

Finally, we applied orthogonal hierarchical clustering to the top driver genes identified across all bacterial networks to visualize their functional polarity at the gene level. The resulting targeted heatmap (Figure 6) displays a striking and sharply resolved dichotomy. Pathogenic organisms—*Porphyromonas gingivalis*, *Prevotella intermedia*, *P. stomatis*, and *Capnocytophaga gingivalis*—show strong activation of the inflammatory–metabolic signature, with genes such as NFKB1, RELA, TNF, MYC, ACACA, and downstream inflammatory mediators consistently elevated. In contrast, commensal species—including *Lactobacillus*, *Streptococcus gordonii*, and *L. plantarum*—exhibit uniformly suppressed activity across these same genes.

**Figure 6.**
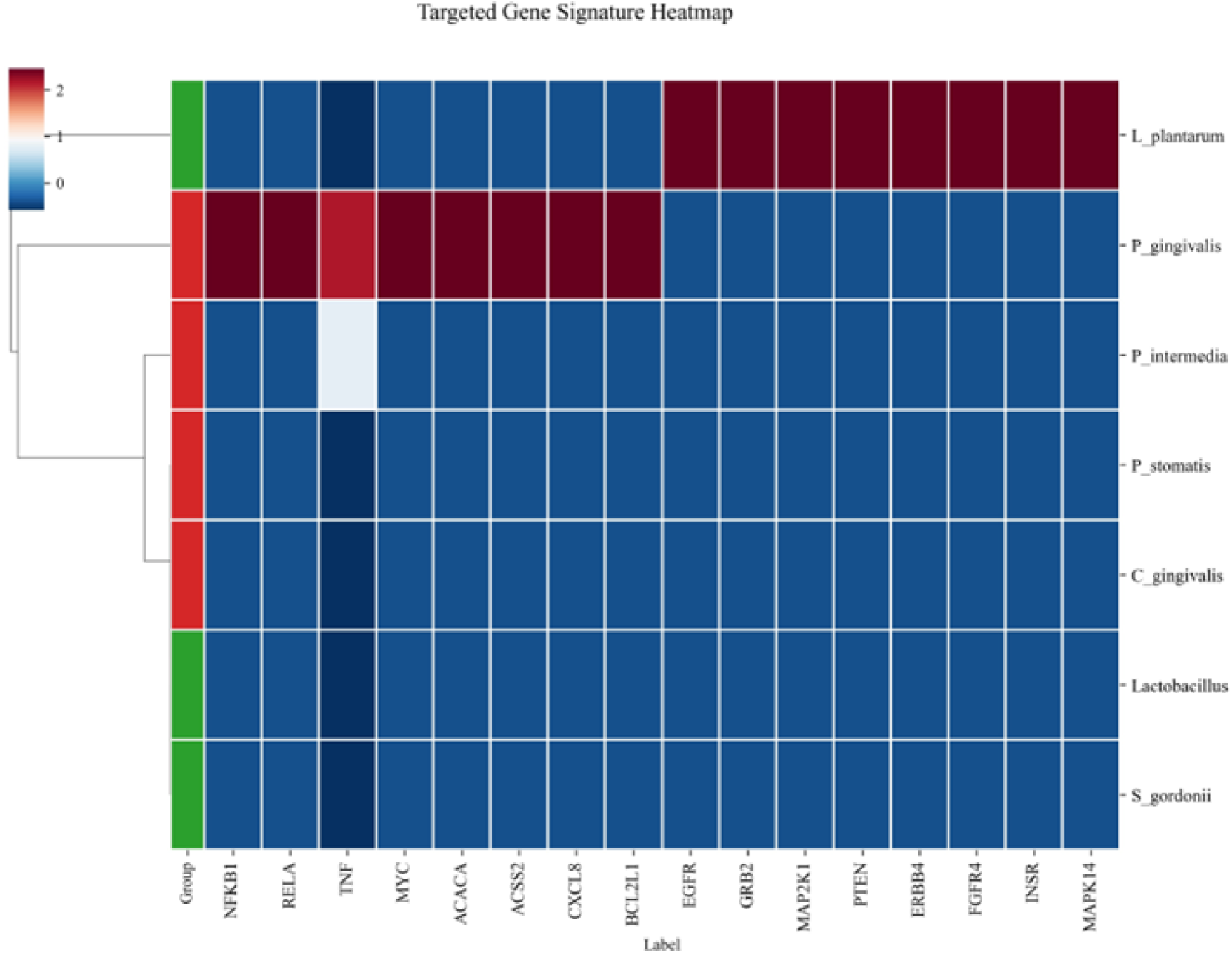
Orthogonal Hierarchical Clustering Reveals Binary Functional Separation of Pathogenic and Commensal Bacteria. Heatmap of the top driver genes involved in inflammatory, metabolic, and kinase-homeostasis signaling across ten oral bacterial species. Red indicates high gene-activity (upregulated/inflammatory) states, and blue indicates suppressed or inactive states. Pathogenic species (*P. gingivalis*, *P. intermedia*, *P. stomatis*, *C. gingivalis*) cluster tightly together, exhibiting strong activation of inflammatory–metabolic regulators (*NFKB1, RELA, TNF, MYC, ACACA*). In contrast, commensal bacteria (*Lactobacillus*, *S. gordonii*) form a distinct, uniformly low-activity cluster. The absence of mixed patterns demonstrates a clean, two-class functional topology, highlighting that inflammatory-metabolic reprogramming is a **unique and exclusive virulence mechanism** of pathogenic taxa.

This “red–blue” separation reflects a binary functional architecture: inflammatory and immunometabolic drivers are exclusively “ON” in pathogens and stably “OFF” in commensals, with no intermediate profiles observed. The clarity of this pattern reinforces that pathogenic bacteria employ a targeted inflammatory-metabolic activation program, whereas beneficial organisms preserve a homeostatic, kinase-regulated epithelial environment.

### 6. Clinical and Translational Validation of Machine Learning–Prioritized Cell Death and Autophagy Networks

To rigorously establish the clinical relevance and translational robustness of our machine learning–derived predictions, we interrogated the expression profiles of prioritized gene sets—encompassing global regulatory hubs, homeostasis-associated signatures, and process-specific markers—across independent clinical transcriptomic and proteomic datasets. Analysis of the GSE37991 cohort revealed marked and statistically significant overexpression of the predicted *Pathogenic Signature* genes in oral squamous cell carcinoma (OSCC) tissues compared with normal oral mucosa. Notably, central regulatory nodes including STAT3, MYC, and CTNNB1, together with the core autophagy adaptor SQSTM1, were robustly upregulated in patient-derived OSCC samples (Figure 7). These findings provide strong clinical confirmation that the microbial-targeted and network-prioritized genes identified by our computational framework are not merely theoretical constructs but represent biologically active and disease-relevant alterations in human OSCC.

**Figure 7.**
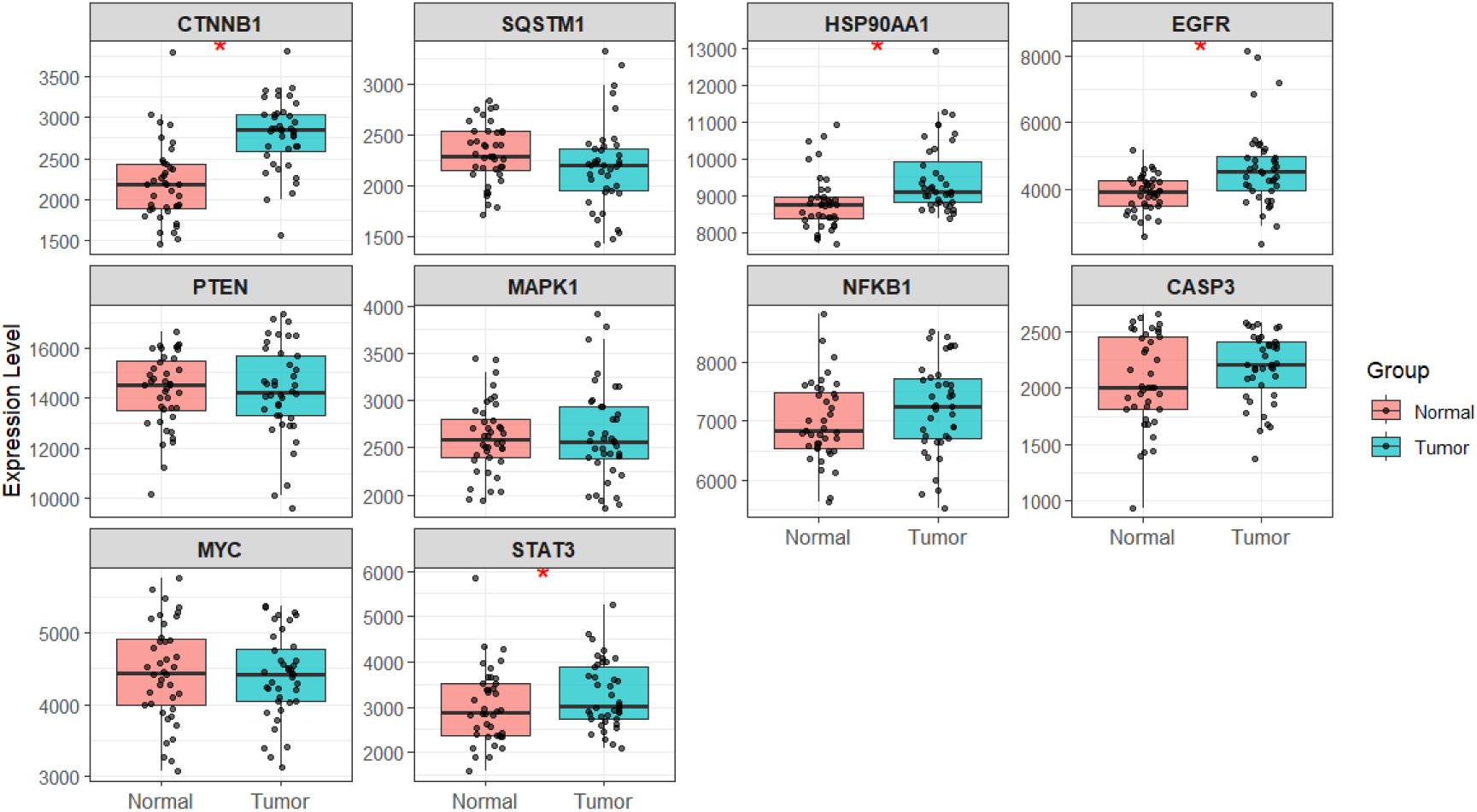
Transcriptomic validation of prioritized autophagy- and cell fate–associated hub genes in OSCC. Box-and-whisker plots illustrate differential mRNA expression of machine-learning–prioritized hub genes (CTNNB1, SQSTM1, HSP90AA1, EGFR, PTEN, MAPK1, NFKB1, CASP3, MYC, and STAT3) in normal oral mucosa versus OSCC tumor tissues, derived from the GSE37991 transcriptomic cohort. Individual data points represent independent patient samples, with box boundaries indicating the interquartile range, the central line denoting the median, and whiskers extending to 1.5× the interquartile range. Red asterisks denote statistically significant differential expression between normal and tumor groups (*p* < 0.05). Notably, key regulatory hubs implicated in autophagy, stress adaptation, and oncogenic signaling, including STAT3, EGFR, HSP90AA1, SQSTM1, and CTNNB1, exhibit marked upregulation in OSCC tissues, supporting their role as clinically relevant nodes within pathogenic cell fate networks. Collectively, these data provide transcriptomic evidence that the computationally identified hub genes correspond to robust molecular alterations in human OSCC, reinforcing the translational validity of the integrative machine-learning framework.

This transcriptomic validation was further reinforced at the protein level using independent immunohistochemical data from the Human Protein Atlas (HPA). Consistent with the RNA-level findings, STAT3 and EGFR exhibited a clear shift from medium staining intensity in normal oral epithelium to high intensity in OSCC specimens, underscoring their activation in malignant tissue. Importantly, HSP90AA1, a key hub within the homeostasis-associated network, was undetectable in normal oral mucosa but displayed medium staining intensity in OSCC, highlighting its selective induction during malignant transformation. While CTNNB1 showed high baseline expression in both normal and tumor tissues, its inclusion as a conserved network hub supports its role as a permissive signaling backbone rather than a binary on/off marker. Collectively, these multi-level validations demonstrate that the computationally identified hubs and signatures translate into bona fide molecular alterations in OSCC patients, thereby strengthening the biological credibility and clinical relevance of our integrative machine learning framework (Figure 8).

**Figure 8.**
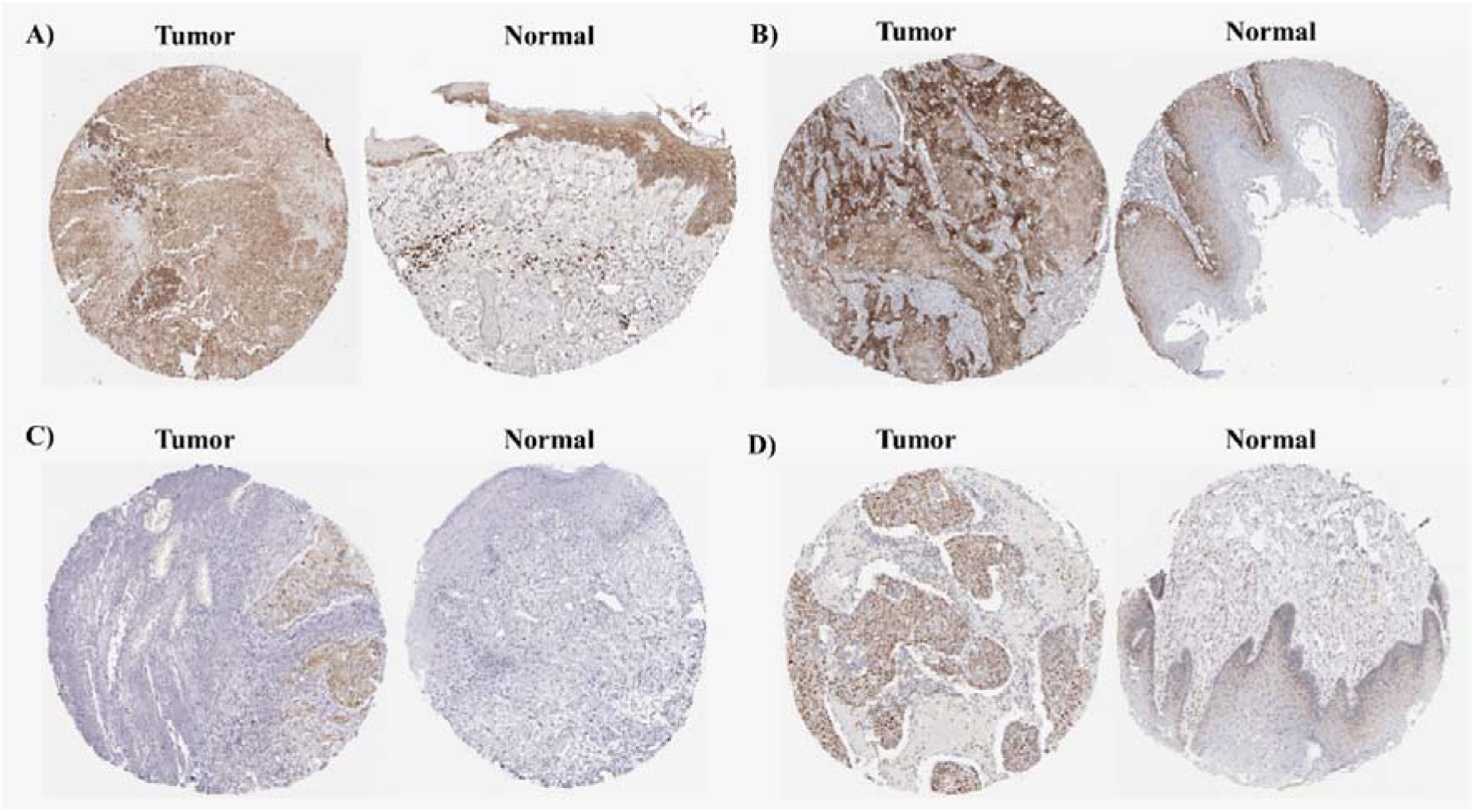
Immunohistochemical validation of machine-learning–prioritized autophagy and cell fate hub proteins in OSCC. Representative immunohistochemical (IHC) staining images from the Human Protein Atlas (HPA) illustrate differential protein expression in OSCC tumor tissues compared with normal oral mucosa for (A) CTNNB1 (Antibody HPA029159; both tumor and normal specimens from female donors), (B) EGFR (Antibody CAB080475; tumor: female, normal: male), (C) HSP90AA1 (Antibody CAB002058; tumor: female, normal: male), and (D) STAT3 (Antibody CAB003859; tumor: female, normal: male). Tumor specimens display markedly increased staining intensity and broader distribution of these proteins relative to normal tissues, indicating activation of key regulatory hubs involved in autophagy, stress adaptation, and oncogenic signaling.

Specifically, EGFR and STAT3 show a clear transition from low-to-moderate staining in normal oral epithelium to strong and widespread immunoreactivity in OSCC, consistent with enhanced proliferative and survival signaling. HSP90AA1, minimally detected in normal tissues, exhibits distinct induction in tumor sections, highlighting its role in malignant transformation and proteostasis remodeling. CTNNB1 demonstrates sustained high expression across tissues, supporting its function as a conserved signaling backbone rather than a binary tumor marker. Collectively, these protein-level data corroborate the transcriptomic findings and confirm that the computationally identified hubs correspond to clinically manifest molecular alterations in OSCC, reinforcing the translational relevance of the machine-learning framework.

## Discussion

Our integrative interactome analysis uncovers two distinct bacterial states within the oral squamous cell carcinoma microenvironment: tumor-promoting (pathogenic) and tumor-restraining (commensal). These states differ fundamentally in how they reprogram autophagy and its crosstalk with EMT, apoptosis, and metabolism. This supports growing evidence that the oral Bacterium is not a passive entity but an active modulator of oncogenic stress adaptation in oral cancer [24–26]. We propose that the bacterial composition critically shapes whether autophagy reinforces invasive plasticity or sustains cytoprotective homeostasis.

Pathogenic taxa—including *Capnocytophaga gingivalis*, *Fusobacterium nucleatum*, *Porphyromonas gingivalis*, *Peptostreptococcus stomatis*, and *Prevotella intermedia*—display conserved signalling strategies where autophagy supports epithelial remodeling and survival. *P. gingivalis* promotes EMT through a MAPK1-driven axis, linking autophagy to invasive phenotype acquisition [19]. In OSCC, PI3K/AKT/mTOR and BCL2–Beclin-1 interactions are known to regulate autophagy–apoptosis balance, supporting survival and adaptability under cellular stress [2]. *F. nucleatum* enhances OSCC proliferation and invasiveness through β-catenin and matrix metalloproteinase pathways [27]. Both *P. gingivalis* and *P. stomatis* activate a MAPK1–STAT3–SRC tri-hub that orchestrates cytokine signaling and cytoskeletal remodeling. Chronic *P. gingivalis* exposure leads to STAT3 phosphorylation and EMT-like shifts in oral epithelia [28, 29], while c-Src has been implicated in actin turnover and EMT across OSCC and head and neck squamous cell carcinoma. Its inhibition impairs motility and invasion, further validating the SRC component [30–33].

*Prevotella intermedia*, frequently enriched in OSCC, drives tumor-promoting inflammation through IL-17 and IL-6 activation, suggesting pyroptosis-autophagy interplay [34]. Pyroptotic pathways, including CASP3–dependent circuits, have been implicated in tumor remodeling, supporting the functional convergence of autophagy and inflammatory death in OSCC [35, 36]. Additionally, *P. gingivalis* outer-membrane vesicles foster proliferation and autophagy activation in epithelial cells and have been shown to manipulate mitophagy in airway tissues [37]. These data support a model in which pathogens exploit autophagy for cytoskeletal reorganization, stress tolerance, and immune escape.

In contrast, probiotic and commensal bacteria such as *Lactiplantibacillus plantarum*, *Lactobacillus spp*., *Streptococcus gordonii*, and *Streptococcus mitis* guide autophagy toward redox homeostasis and epithelial resilience. *S. anginosus* appears intermediate, maintaining low-grade inflammatory signaling without overt invasive signatures. *L. plantarum* fosters a MAPK-suppressive and PTEN-activating program, promoting apoptosis and limiting proliferation in OSCC models [38]. Similarly, *L. acidophilus* exhibits anti-proliferative properties and induces apoptosis, supporting the tumor-restraining effect of Lactobacillus species [39]. *L. plantarum*-derived postbiotics have also demonstrated anticancer potential, including chemosensitization [40]. Although direct experimental validation of the MAPK1–STAT3–HSP90AA1–LDHA/TKT axis in Lactobacillus-treated OSCC is limited, prior studies suggest this genus influences MAPK/PI3K–AKT signaling and oxidative stress responses in epithelial contexts [41].

*Streptococcus gordonii* showed transcriptional enrichment in detoxification and inflammatory regulation, aligning with its ability to protect epithelial barriers and inhibit pathogenic colonization by *F. nucleatum* [42]. S. mitis supports redox balance through activation of the AhR pathway, thereby enhancing xenobiotic metabolism and antioxidant signaling in oral epithelial cells [43]. Collectively, commensal Streptococci regulate MAPK, NF-κB, and detoxification pathways, steering autophagy toward protective cellular programs that counteract pro-invasive stress.

We identify three autophagy-centric signaling architectures that may determine tumor-promoting versus tumor-suppressive outcomes in OSCC: (1) a MAPK1–STAT3–SRC inflammatory-invasive circuit; (2) a MAPK1–STAT3–CASP3–GSDMD pyroptotic-inflammation module; and (3) a MAPK1–HSP90AA1/CTNNB1/GSK3B proteostasis-centered homeostatic scaffold [44, 45]. Metabolic subnetworks—including LDHA/TKT and antioxidant/xenobiotic mediators like GSTP, ALDH, CAT, SOD1, and NFKB1—likely modulate the tipping point between cytotoxic and regenerative autophagy. This gradient mirrors bacterium signatures in OSCC: *F. nucleatum* and *P. intermedia* correlate with aggressive clinical features and progression [46], while probiotic *Lactobacilli* restrict OSCC cell proliferation and support apoptosis, offering prognostic and therapeutic promise [39].

Therapeutically, our findings highlight MAPK1, STAT3, SRC, PTGS2, and GSDMD as candidate nodes for targeted modulation. Pharmacologic inhibition of STAT3 has been shown to reduce OSCC cell proliferation and EMT [47], while COX-2 blockade restores epithelial adhesion and reverses invasive phenotypes. Early trials suggest that bacterium reprogramming—via probiotic combinations may favorably alter the tumor-immune axis and enhance chemosensitivity [48]. *S. mitis* also modulates epithelial repair through AhR activation, further supporting its protective profile [43].

Machine learning added critical structural validation. Our hierarchical PCA and pathway scoring resolved dominant and secondary topological patterns, aligning bacterial identity with network-level host impacts. *F. nucleatum* and *S. mitis* emerged as orthogonal extremes—one anchoring survival-autophagy programs and the other mapping onto detoxification-autophagy axes. Removal of these outliers exposed secondary programs, such as inflammatory-metabolic (*P. gingivalis*) and kinase-homeostatic (*L. plantarum*) signatures. The empirical p-value (p < 0.001) from permutation testing confirmed non-randomness, while linear discriminant analysis and the Dysbiosis Index demonstrated that these topologies encode stable, biologically coherent stratifications.

Clinical validation using independent microarray datasets and the Human Protein Atlas provides a strong translational link between our computational framework and human OSCC pathology. The concordant upregulation of the global hubs STAT3, MYC, and CTNNB1 at both mRNA and protein levels confirms our model’s prediction that pathogenic microbial signatures converge on these nodes as central regulators of oncogenic transformation. Consistent with this, STAT3 activation is a hallmark of OSCC driven by inflammatory cues within the microbe-enriched tumor microenvironment [44], while MYC overexpression promotes metabolic reprogramming and rapid tumor cell proliferation [49].

We further observed induction of HSP90AA1, transitioning from “not detected” in normal tissues to “medium intensity” in OSCC, supporting its role as a master chaperone stabilizing oncogenic kinases and sustaining pathogenic signaling networks [50]. Elevated SQSTM1 expression is consistent with an aberrant autophagic state, where p62 accumulation may reflect impaired autophagic flux and enhanced NRF2-mediated stress adaptation, fostering tumor cell survival [51]. Although EGFR and STAT3 show stronger tumor-specific staining, the persistently high expression of CTNNB1 across both tissue types suggests regulation at a functional level, such as pathogen-induced nuclear translocation, rather than expression abundance [52]. Collectively, these multilayered data demonstrate that OSCC exhibits clinically measurable molecular rewiring captured by both the Dysbiosis Index and the machine learning–derived network topology, reinforcing the translational relevance of our integrative approach.

Together, these findings indicate that topological differences in bacterial host-network embedding reflect functionally encoded divergence. Pathogens exploit survival-autophagy plasticity; commensals maintain cytoprotective fidelity.

## Conclusion and Future Directions

Our findings reposition the oral microbiome as an active regulatory axis—a molecular switchboard that governs the balance between autophagy-driven invasion and autophagy-mediated epithelial preservation in OSCC. Rather than acting as passive bystanders, oral bacteria encode pro-tumorigenic or cytoprotective signals within the topology of host–microbe interaction networks, thereby determining whether autophagy is co-opted to support inflammation and survival or deployed to maintain epithelial homeostasis. This network-level encoding of biological intent offers actionable leverage points for both risk stratification and therapeutic intervention.

Importantly, these results argue that OSCC risk may be modifiable through targeted reshaping of oral microbial community structure. Shifts in the relative abundance of pathogenic versus commensal taxa—driven by diet, oral hygiene, antibiotics, inflammation, or aging—are predicted to recalibrate autophagy polarity at the epithelial interface. Future studies should therefore test whether restoring commensal dominance, suppressing pathogenic network drivers, or stabilizing microbial community resilience can actively reduce OSCC susceptibility, delay malignant progression, or enhance therapeutic responsiveness.

While this study focused on bacterial–host interactions, the contributions of oral fungi and viruses to OSCC-associated autophagy rewiring remain unexplored and warrant systematic investigation. Future work should prioritize functional perturbation of key network nodes identified here—STAT3, SRC, PTGS2, GSDMD, HSP90AA1, GSK3B, and CTNNB1—under defined microbial exposures. Experimental platforms integrating CRISPR-based pathway modulation, epithelial organoid–microbe co-cultures, and spatially resolved multi-omics will be essential to dissect causality, directionality, and context dependence within bacterium–epithelium signaling circuits.

In parallel, integrative metabolomics–autophagy profiling will be critical for identifying microbial metabolites (e.g., short-chain fatty acids, quinolones, and amino acid derivatives) that converge on Ca²□/MAPK/mTOR signaling axes. Of particular interest is chemosensory regulation: bitter (T2R) and sweet (T1R) taste receptors expressed by oral epithelia function as microbial metabolite sensors and modulators of MAPK–mTOR–STAT3 pathways. We hypothesize that T2R activation promotes autophagy-linked epithelial protection, whereas T1R signaling biases toward inflammatory survival programs. Pharmacologic or genetic manipulation of these receptors may therefore represent a previously underappreciated strategy to reprogram bacterium–autophagy crosstalk.

Ultimately, rational manipulation of oral microbial ecosystems—through diet, prebiotics, probiotics, postbiotics, or engineered bacterial consortia such as *Lactobacillus plantarum* combined with *Streptococcus mitis*—may offer a feasible route to restore autophagy homeostasis and suppress tumor-permissive signaling. Together, these findings establish a conceptual and experimental framework for precision microbiome-guided prevention and therapy in oral oncology, positioning microbial community engineering as a viable strategy to reduce OSCC risk and alter disease trajectory.

## Author Contributions

**Hamid Latifi Navid** and **Rui Vitorino** contributed to **data mining, systems-level network construction, and integrative network analysis**. **Mahdi Akhavan** and **Iman Beheshti** performed the **machine learning analyses**, including model development, coding, optimization, and methodological implementation. **Pooya Jalali** conducted **independent validation analyses** to confirm the robustness and reproducibility of the computational findings. **Amir Barzegar Behrooz** and **Sanaz Vakili** contributed to **manuscript drafting, figure development, and integration of computational and biological interpretations**. **Anil Menon**, **Vimi S. Mutalik**, and **Robert J. Schroth** served as **oral health and clinical advisors**, providing domain expertise, clinical interpretation, and guidance on translational relevance. **Prashen Chelikani** contributed to **conceptual consolidation and interdisciplinary synthesis** of the study. **Saeid Ghavami** conceived the study, **developed the central hypothesis**, led and coordinated the research team, supervised all analytical and interpretative components, and performed the **final manuscript compilation and critical revision**. All authors reviewed and approved the final version of the manuscript.

## Acknowledgments

The authors acknowledge the use of ChatGPT 5.2 for limited assistance with English language editing, conducted under full human oversight and responsibility.

